# Genomic and radiocarbon insights into the mystery of mouse mummies on the summits of >6000 m Andean volcanoes

**DOI:** 10.1101/2023.04.22.537927

**Authors:** Jay F. Storz, Schuyler Liphardt, Marcial Quiroga-Carmona, Naim M. Bautista, Juan C. Opazo, Timothy B. Wheeler, Guillermo D’Elía, Jeffrey M. Good

**Author notes:** Correspondence (JFS).

## Abstract

Our understanding of the limits of animal life is continually revised by scientific exploration of extreme environments. Here we report the discovery of numerous mummified cadavers of leaf-eared mice, *Phyllotis vaccarum*, from the summits of three different Andean volcanoes at elevations 6029-6233 m (19,780-20,449 ft) above sea level in the Puna de Atacama (Chile-Argentina). Such extreme elevations were previously assumed to be completely uninhabitable by mammals. In combination with a live-captured specimen of the same species from the nearby summit of Volcán Llullaillaco (6739 m [=22,110 ft]), the 13 summit mummies represent the highest physical records of mammals in the world. We report a chromosome-level genome assembly for *P. vaccarum* in combination with a whole-genome re-sequencing analysis and radiocarbon dating analysis that provide insights into the provenance and antiquity of the summit mice. We test alternative hypotheses to explain the existence of mouse graveyards on the summits of Atacama volcanoes. Radiocarbon data indicate that the most ancient of the mummies were at most a few centuries old. Genomic polymorphism data revealed a high degree of continuity between the summit mice and conspecifics from lower elevations in the surrounding Altiplano. Genomic data also revealed equal numbers of males and females among the summit mice and evidence of close kinship between some individuals from the same summit groups. These findings bolster evidence for self-sustaining populations of *Phyllotis* at elevations >6000 m and challenge assumptions about the environmental limits of vertebrate life and the physiological tolerances of small mammals.

## INTRODUCTION

“Kilimanjaro is a snow covered mountain 19,710 feet high and is said to be the highest mountain in Africa…Close to its summit there is the dried and frozen carcass of a leopard. No one has explained what the leopard was seeking at that altitude.”

-*The Snows of Kilimanjaro*, Ernest Hemingway

Exploration of extreme environments such as deep ocean trenches, the Antarctic sea floor, and high mountain summits have led to discoveries of animal life flourishing in unexpected and surprisingly hostile conditions^1-3^. In a recent mountaineering mammal survey of Volcán Llullaillaco, a stratovolcano in the Central Andes that straddles the border between Argentina and Chile, numerous specimens of the Andean leaf-eared mouse *Phyllotis vaccarum* (previously known as *Phyllotis xanthopygus rupestris*) were captured at or above the elevational limits of vegetation, >5000 m above sea level, and one specimen was captured on the very summit of the volcano at 6739 m (22,110 ft)^4^. On the summits of >6000 m Andean volcanoes like Llullaillaco, plant life is nonexistent, temperatures less than -30°C are common even in summertime, wind speeds can reach 200 km/hr, and there is less than half the oxygen available at sea level. For such reasons, these extreme elevations were previously thought to be completely uninhabitable by mammals. It remains unclear whether records of mice in such hostile environments reflect rare dispersal events by transient sojourners, recent upslope colonization facilitated by climate change, or long-term persistence of resident populations.

Here we report the discovery of mummified cadavers of *Phyllotis* on the summits of three different >6000 m volcanoes in the Puna de Atacama, Central Andes, including concentrated aggregations of multiple mummies on the same summits. Along with the above-mentioned summit mouse from Llullaillaco, these vouchered mummy specimens represent the highest physical records of mammals in the world. We report whole-genome sequence data and radiocarbon data that provide insights into the colonization history and antiquity of the *Phyllotis* mummies.

The first published reports of mice at elevations >6000 m came not from biologists but from archaeologists, as mouse mummies have occasionally been discovered in association with Incan ceremonial structures and burial sites at or near the summits of several high Andean volcanoes^5,6^. However, it was always assumed that the mice had been transported to such sites by Incas, either inadvertently or because the animals were used as part of a sacrificial ritual. Le Paige^5^ (pp. 40-41) describes the discovery of mouse mummies in association with an Incan ceremonial structure on the summit of Volcán Púlar (6233 m) in northern Chile [translated from Spanish]: “We were surprised to find two mice on the top of this volcano, one of them inside the ceremonial structure and the other immediately outside it. Apparently, these rodents arrived here in bundles of firewood…or they could have been part of a ritual…”. Similarly, mummified mice were discovered in association with Incan ceremonial structures on the summits of Cerro El Toro (6168 m) and Cerro Las Tόrtolas (6160 m), both on the Argentina-Chile border^6^. Affirming the conventional wisdom among archaeologists, Beorchia^6^ (p. 241) stated [translated from Spanish]: “A rodent does not live at more than 6000 meters because it would have no source of food. Did they come in loads of firewood and grass? It is a somewhat difficult hypothesis to accept … I am inclined to the idea that they were transported intentionally as part of a sacrificial ritual, or for some magical purpose beyond my understanding.”

Quite understandably, the archaeologists quoted above did not consider the possibility that the mice naturally occurred at those extreme elevations and that they had reached the summits of their own accord. In light of the recent live-capture record of a mouse from the 6739 m summit of nearby Volcán Llullaillaco (6739 m)^4^, it appears that mice do in fact live at such elevations, and – when they meet their end – their cadavers, being well preserved by the cold, dry conditions, may occasionally (and coincidentally) be found in association with Incan structures.

Ritualized child sacrifices and related ceremonies (‘Qhapaq hucha’) that were performed on the summits of sacred peaks were conducted by Incas who traveled in caravans from the heart of the empire in Cuzco, Perú, and biological artifacts that were interred with sacrificial victims were typically derived from localities far from the ceremonial sites. For example, child mummies from a burial site below the summit of Llullaillaco, located in the exceedingly arid, desolate terrain of the Puna de Atacama, were adorned with headdresses made of feathers from Amazonian birds and jewelry fashioned from oyster shells from the Pacific Coast of Ecuador^7,8^. Thus, mice or other animals that were used as part of a ritualized mountaintop sacrifice could have geographic origins far distant from their final resting place. In the case of *Phyllotis* from the Andean Altiplano and adjoining regions of South America, geographic patterns of mitochondrial DNA variation have been well-characterized within and among species ^4,9-12^, so genomic sequence data should reveal whether summit mummies were derived from local source populations or whether they represent phylogeographic anomalies that trace their origins to more distant corners of the former Incan empire. Radiocarbon dating permits an even more conclusive test of the Incan transport hypothesis, since the Incan presence in this region is thought to have begun sometime after 1470, lasting until the initial years of the Spanish conquest in 1532 ^7,13^. If the mouse mummies are less than ∼500 yrs old, that would indicate that they were not contemporaries with the Incas. In addition to testing the Incan transport hypothesis, our main motivation for estimating the ages and genetic affinities of the summit mummies was to assess evidence for long-term, self-sustaining populations of mice at elevations >6000 m and to obtain insights into their colonization history.

## RESULTS AND DISCUSSION

### Discovery of summit mummies

In 2020-2022, we conducted mountaineering mammal surveys of 21 volcanoes with elevations 5706-6893 m above sea level in the Central Andes (Figure S1). In addition to live-trapping records of mice from a range of elevations surrounding each volcano, we discovered a total of 13 mouse mummies on the summits of three separate >6000 m peaks: Volcán Salín (6029 m [19,780’]; Argentina-Chile), Volcán Púlar (6233 m [20,449’]; the highest peak in Chile that does not share a border with Argentina or Bolivia), and Volcán Copiapó (6052 m [19,856’], Chile)(Figure 1). On the summits of Salín and Púlar, we excavated mummified cadavers of multiple individual mice from the crevices of volcanic rock piles and alcoves (Figure 2, Figure S2), and in each case the mummified cadavers were found in association with skeletal remains of numerous other mice. Of the four specimens from the summit of Volcán Púlar, one was an intact mummy and the remaining three samples represented partial skeletons. Based on examination of cranial and external characters, we provisionally identified the mummy specimens as leaf-eared mice in the genus *Phyllotis*.

**Figure 1.**
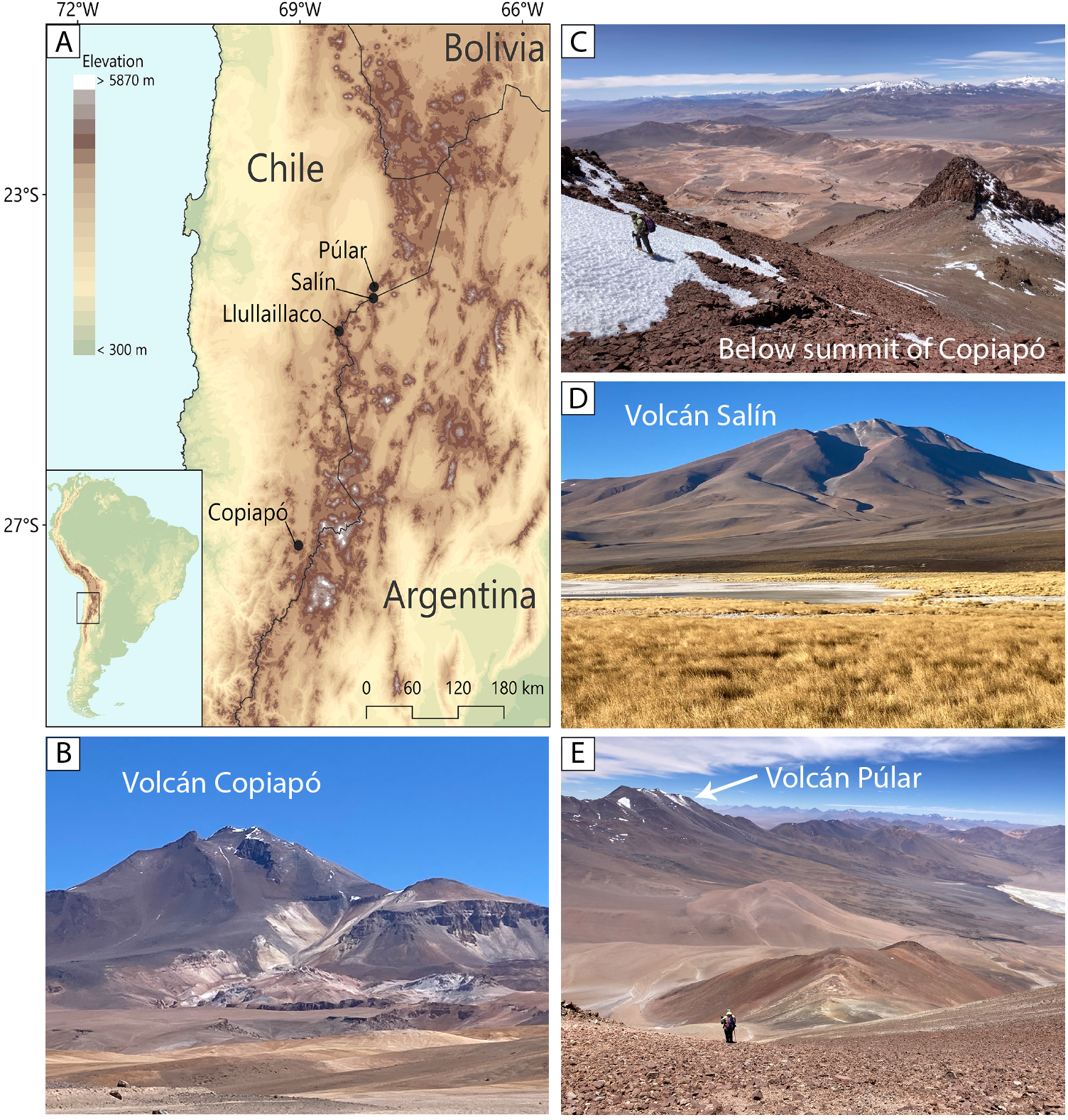
Records of *Phyllotis* mice from summits of four >6000 m volcanoes in the Central Andes. (A) Volcano summits where *Phyllotis* mice were either live-trapped (Llullaillaco)^4^ or discovered as mummies (Púlar, Salín, and Copiapό). (B) Volcán Copiapό (6052 m), Chile, viewed from the North. (C) View to the North from just below the summit of Volcán Copiapό. The right-angle corner of an Incan ceremonial platform can be seen in the foreground, highlighted by red dashed line. (D) Volcán Salín (6029 m), Argentina-Chile, viewed from the North. (E) Volcán Púlar (6233 m), Chile, viewed from the northern flank of Volcán Salín.

**Figure 2.**
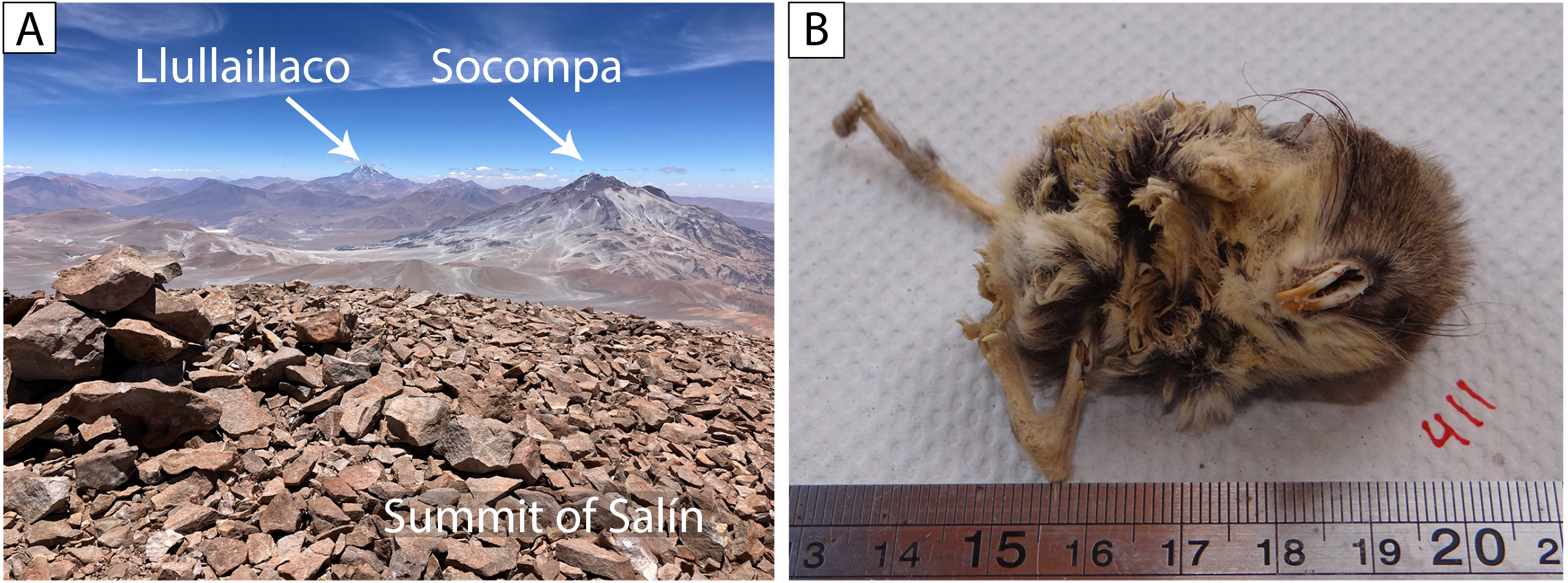
Excavation of *Phyllotis* mummies from volcano summits. (A) Rock pile on the summit of Volcán Salín (6029 m) from which a total of 8 individually intact *Phyllotis* mummies were excavated. Volcán Llullaillaco (6739 m) and Volcán Socompa (6051 m), two other volcanoes with summit records of *Phyllotis*^4,25^, are visible to the South. (B) Representative *Phyllotis* mummy (UACH8545) excavated from the summit of Volcán Salín.

### Antiquity of mummies estimated by radiocarbon dating

Radiocarbon dating revealed that the summit mummies were not contemporaneous with the Incas, as estimated ages were uniformly <500 years BP (Table 1). The Salín and Copiapó specimens were at most a few decades old, as lower bounds of the 95.4% high probability density range are uniformly >1955 AD. The Volcán Púlar samples were at most ∼350 years old, as the lower bounds of the 95.4% high probability density range were 1670-1675 AD and upper bounds were >1950 AD.

**Table 1.**
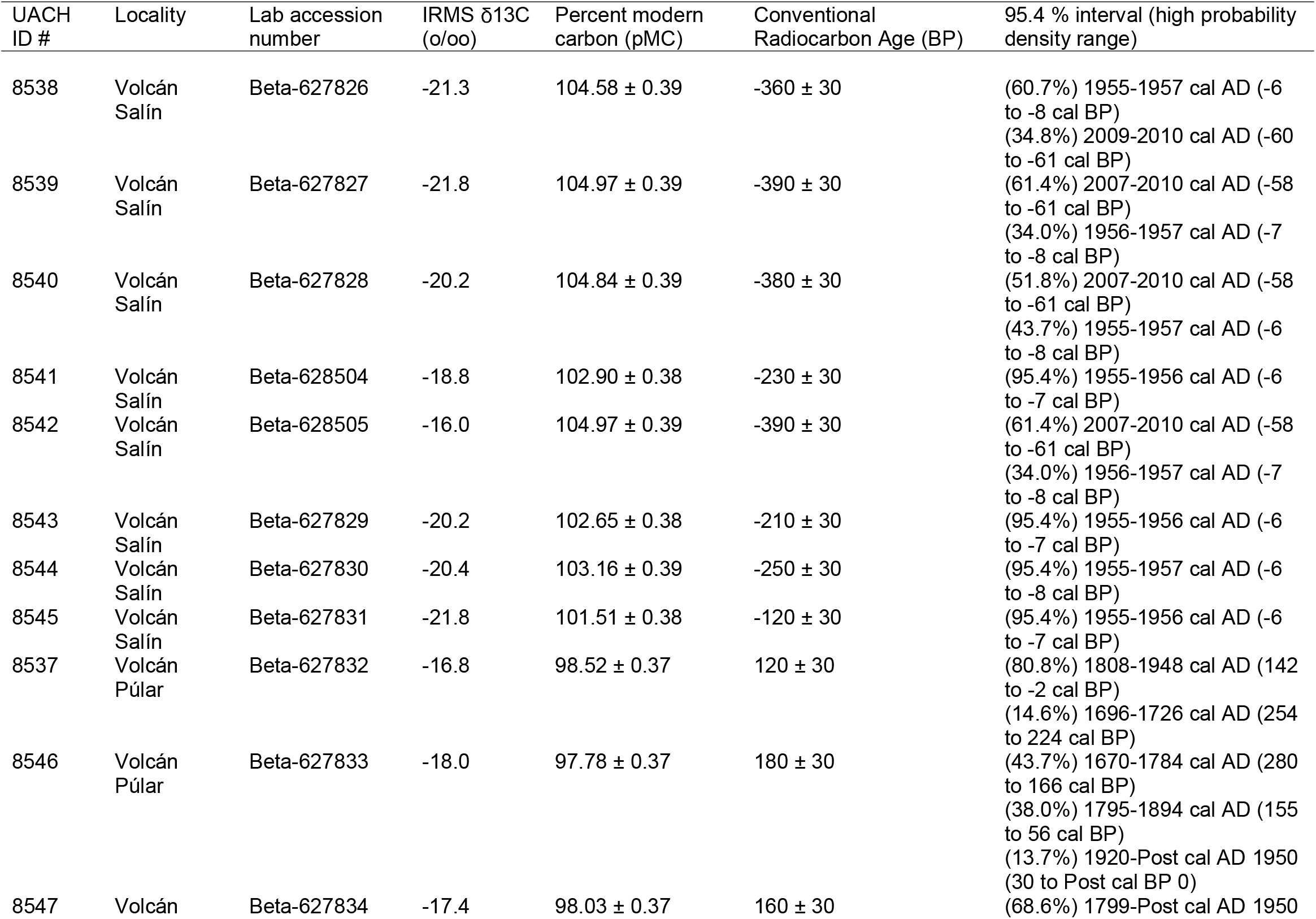
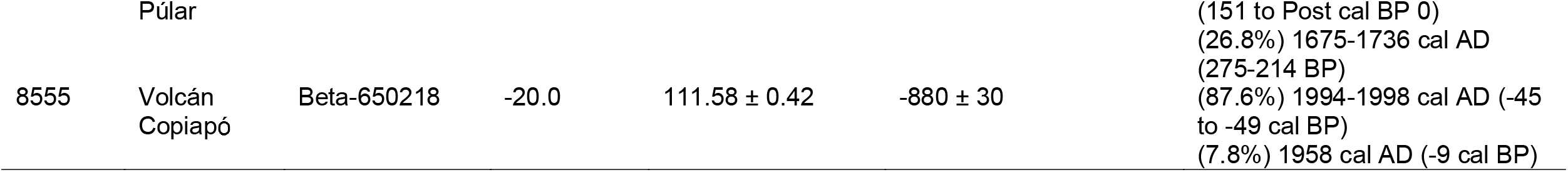
Radiocarbon summary statistics and calibrated date estimates for *Phyllotis* mummies from the summits of Volcán Salín (6029 m), Volcán Púlar (6233 m), and Volcán Copiapό (6052 m), Puna de Atacama, Central Andes. Conventional radiocarbon ages (rounded to the nearest 10 yrs) were used as input for calendar calibration using the SHCAL13+SHZ_2 and SHCAL databases. Errors are reported as 1σ counting statistics; calculated values <30 BP are conservatively rounded up to 30. The estimated percent modern carbon of all specimens from Volcán Salín and Volcán Copiapό were >100 (where the modern reference is 1950 AD), and lower bounds of the 95.4% high probability density range are uniformly >1955 AD, meaning that the samples are at most several decades old. The Volcán Púlar samples are at most ∼350 years old, as the lower bounds of the 95.4% high probability density range are 1670-1675 AD and upper bounds are >1950 AD.

### Assembly of a *de novo* reference genome for *Phyllotis vaccarum*

We used whole-genome sequence data to infer the species identity of the mummies, to infer their geographic provenance, and to gain insight into the colonization history of the summit mice. To date, genetic analyses of *Phyllotis* mice have been restricted to mitochondrial DNA sequence data and the most closely related species with a sequenced reference genome is the Hispid cotton rat, *Sigmodon hispidus*, a species that diverged from *Phyllotis* >10 MYA ^14-18^. We therefore sequenced and assembled a chromosome-level *de novo* reference genome of *P. vaccarum*. We assembled PacBio HiFi long-read sequences (26.7x coverage) followed by chromosome-scale scaffolding using Illumina sequencing of long-range proximity ligation of endogenous chromatin (Omni-C technology). We used these data to generate a 2.66 GB assembly that was highly contiguous (scaffold N50=147.8Mb; N90=59.3Mb) with near complete coverage of conserved eukaryotic orthologs (251 of 255 complete BUSCO copies, 98.4%). Most of the genome was assembled into 19 large scaffolds (Figure S3), consistent with a complete chromosome-scale assembly matching the described karyotype for the species (2N=38)^19^.

### Population re-sequencing

We next generated low coverage whole-genome sequences of 41 mice (0.8-12.2x coverage, median 2.3x), including 13 mummies from Volcán Salín (*n*=8), Volcán Púlar (*n*=4), and Volcán Copiapó (*n*=1), the single live-trapped mouse from the summit of Llullaillaco, and 27 additional live-trapped specimens from lower elevation sites (3651-5070 m) in the region surrounding the afore-mentioned volcanoes (Figure 3A). Using ancient DNA extraction protocols, we successfully generated high-quality sequence data for all mummies and skeletal samples. Volcán Salín mummies showed no appreciable environmental contamination with >99% of reads mapping for all samples (Table S1). Between 26.0-81.5% of all sequence reads from the mummies and skeletal material from Púlar and Copiapó derived from endogenous mouse DNA, with the vast majority of contamination coming from microbial sequences. Analysis of nucleotide misincorporation profiles revealed no evidence of elevated DNA damage that is typical of ancient or historical samples ^20-22^, suggesting that the sequence quality of the summit mummies was equivalent to that of contemporary material (as indicated by radiocarbon data for the and Salín and Copiapó mummies; Table 1). If the Púlar samples were on the order of several centuries old, as suggested by the upper range of radiocarbon dating estimates (Table 1), they were clearly well-preserved in the perpetually freezing and extraordinarily arid conditions that prevail on the summits of >6000 m Atacama volcanoes.

**Figure 3.**
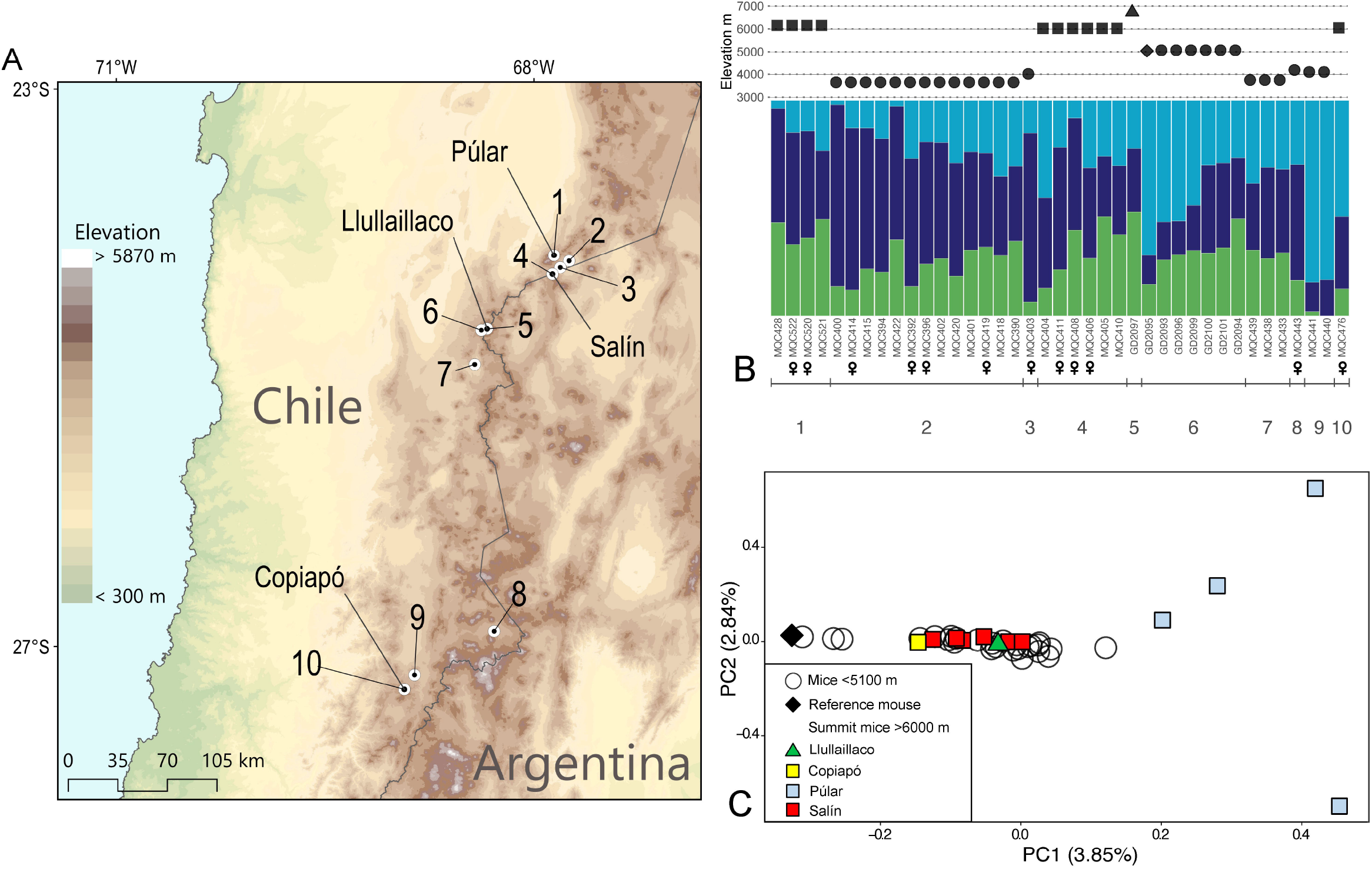
Geographic sampling and the patterning of genomic variation in *Phyllotis vaccarum* from the summits, flanks, and surrounding regions of four >6000 m volcanoes: Púlar, Salín, Llullaillaco, and Copiapό. (A) Sampling localities for mice from >6000 m summits and conspecifics from lower elevations in the surrounding Altiplano. (B) Structure analysis reveals low levels of population structure across the surveyed region, as indicated by similar estimates of admixture proportions in 37 specimens of *Phyllotis vaccarum* from spatially disparate sites that span a range of elevations (3651-6739 m). Samples are grouped by locality and numbered according to panel A. Female specimens are denoted by ‘♀’. The elevation at which each specimen was collected is displayed above the estimated admixture proportions; squares denote summit mummies, the triangle denotes the live-caught mouse from the summit of Llullaillaco, the diamond denotes the mouse that served as the source of the reference genome, and circles denote live-caught mice from the surrounding Altiplano. (C) Principal component analysis (PCA) of genomic variation in the same set of samples shown in panel B.

Our sample of 14 mice from >6000 m volcano summits included an equal number of males and females (the live-captured Llullaillaco mouse was male, and genomic analysis revealed that the set of summit mummies included 6 males and 7 females). The sample of summit mice from Púlar included 2 males and 2 females, and the sample from Volcán Salín included 4 males and 4 females. The Salín sample also included two pairs of close relatives (coefficient of kinship, Θ = 0.427, 0.418), representing full-siblings or parent-offspring pairs, whereas remaining individuals were not closely related (Θ < 0.01). Since dispersal in small rodents is typically sex-biased and generally occurs over short distances^23^, the equal representation of males and females in our sample of summit mice and evidence for close kinship within groups from the same summit is consistent with the idea that the summit mice represent members of self-sustaining populations and were not simply transient sojourners.

Average genome-wide pairwise nucleotide diversity for the total sample of mice was typical for natural populations of rodents (π = 0.42%)^24^. After filtering our dataset to exclude closely related individuals, we confirmed that the summit mice exhibited close affinities to the *Phyllotis vaccarum* reference genome and other live-trapped specimens of *P. vaccarum* from lower elevations (3651-5070 m) on the flanks of the same volcanoes and in the surrounding Altiplano (Figure 3B, C). Results of a model-based clustering analysis revealed very low population structure in the total sample, as proportional assignments of ancestry to genetically defined clusters were similar for mice from all localities (Figure 3B, Figure S4). A principal component analysis of genomic variation also revealed low levels of population structure across the surveyed region, as PC1 and PC2 explained only 4.05% and 3.00% of total variation, respectively (Figure 3C). The patterning of genomic variation did not distinguish the set of summit mice from mice collected at less extreme elevations in the same region, although the four Volcán Púlar mice were somewhat distinct from each other and they clustered apart from mice from the other localities (Figure 3C). This pattern could reflect a temporal dimension of allele frequency variation given that radiocarbon dating suggested that those four samples could be several centuries old and may therefore be separated in time from one another and from the contemporary specimens by several hundreds of mouse generations.

### New elevational records for wild mammals

In combination with the live-captured specimen of *Phyllotis vaccarum* from the summit of Volcán Llullaillaco (6739 m)^4^, the vouchered summit mummies from 6029-6233 m represent the highest elevation mammal specimens on record. Video records, evidence of active burrows, and various lines of indirect evidence indicate that *Phyllotis* mice occur at elevations >6000 m on Llullaillaco and neighboring Volcán Socompa^4,25,26^. Thus, the highest records of *Phyllotis* (and the only physical records of wild mammals from elevations >6000 m) are all from the summits of Andean peaks within a section of the Central Volcanic Zone in the Puna de Atacama, between latitudes of 24°S and 28°S, South of the Tropic of Capricorn.

## Conclusions

The summit mummies were not found in close association with Incan ceremonial structures and radiocarbon data confirm that the mummies are at most a few centuries old (in the case of the Púlar specimens) or no more than a few decades old (in the case of the Salín and Copiapό specimens). It is clearly not necessary to invoke transport by Incas to explain the existence of dead mice on the summits of peaks like Púlar and Copiapό where ceremonial structures have been discovered ^5,6^. Moreover, population genomic data indicate that the summit mice are representative of the regional population in the surrounding Altiplano; they clearly do not derive from a distinct source population. These findings, in conjunction with the observation of equal numbers of males and females in the sample of summit mice and evidence of close kinship between some mice from the same summit groups, bolster evidence for long-term resident populations of *Phyllotis* at elevations >6000 m.

It is unclear why the mice are found in concentrated aggregations in summit graveyards, and their reasons for ascending to such heights in the first place are equally inscrutable. Like the Kilimanjaro leopard mentioned by Hemingway, we have no explanation for what the animals were seeking at those altitudes. Mice climb to the icy, wind-scoured summits of >6000 m Andean volcanoes for reasons that remain mysterious – perhaps just as mysterious as why the Incas chose to ascend the same summits to perform ritualized child sacrifices.

## ACKNOWLEDGMENTS

This work was funded by grants from the National Institutes of Health (R01 HL159061, JFS and JMG), National Science Foundation (IOS-2114465, JFS; OIA-1736249, JFS and JMG), National Geographic Society (NGS-68495R-20, JFS) and the Fondo Nacional de Desarrollo Científico y Tecnológico (Fondecyt 1221115, GD). We thank Mario Pérez-Mamani, Juan Carlos Briceño, and Alex Damian González Sandoval for assistance and companionship in the field, Brandi Coyner (Sam Noble Museum of Natural History, University of Oklahoma) for frozen tissue loans, members of the Good lab and the UNVEIL network for helpful discussions, and the University of Montana Genomics Core for access to instrumentation. We also acknowledge the assistance of Dovetail Genomics in producing the reference genome assembly. Computational resources were provided by the Griz Shared Computing Cluster, University of Montana.

## AUTHOR CONTRIBUTIONS

J.F.S., S.L., M.Q-C., G.D., and J.M.G designed the research; J.F.S, M.Q-C., N.M.B, and G.D. performed the field work; S.L. and T.W. performed the laboratory work; S.L., J.C.O., G.D., and J.M.G. analyzed data; J.F.S., S.L., M.Q-C., N.M.B., G.D., and J.M.G. prepared figures and wrote the manuscript.

## METHODS AND MATERIALS

### Data and code availability

- All genome sequences and the reference genome were deposited in GenBank under BioProject PRJNA950396 and are publicly available as of the date of publication.
- This paper does not report original code.
- Any additional information required to reanalyze the data reported in this paper is available from the lead contact upon request.

### Live-trapping and specimen preparation

We live-trapped mice using Sherman traps. We sacrificed mice in the field, prepared them as museum specimens, and preserved tissue samples as sources of DNA. Specimens are housed in the Colección de Mamíferos, Universidad Austral de Chile, Valdivia, Chile. We identified all specimens to the species level based on cranial and external characters^27^, and all such identifications were subsequently confirmed with molecular sequence data.

All mice were collected in accordance with permissions to JFS from the following Chilean government agencies: Servicio Agrícola y Ganadero (SAG, Resolución exenta #’s 6633/2020, 5799/2021, and 3204/2022), Corporación Nacional Forestal (CONAF, Autorización #’s 171219 and 1501221), and Dirección Nacional de Fronteras y Límites del Estado (DIFROL, Autorización de Expedicion Cientifica #68, 02/22, and 07/22). Mice were handled in accordance with protocols approved by the Institutional Animal Care and Use Committee (IACUC) at the University of Nebraska (project ID: 1919).

### Radiocarbon dating

Radiocarbon dating of mummy samples was performed at the Beta Analytic Radiocarbon Dating Laboratory, Miami, FL, using <1 g samples of dried tissue or skeletal material from each individual specimen. Conventional radiocarbon age was calculated using the Libby half-life (5,568 yrs) corrected for total isotopic fraction. The radiocarbon age is rounded to the nearest 10 yrs and is reported as radiocarbon years before present (BP, where ‘present’ = 1950 AD). The conventional radiocarbon ages were used to calibrate calendar years using the SHCAL13 + SHZ1-2 or SHCAL20 databases. d13C values for pre-treated samples were measured separately in an isotope ratio mass spectrometer (IRMS).

### Genome sequencing and assembly

Sequencing and assembly of a *P. vaccarum* reference genome was performed by Dovetail Genomics™ (Santa Cruz, CA). We first sequenced DNA from ethanol-preserved liver tissue from a single live-trapped female *P. vaccarum* from Volcán Llullaillaco (UACH8516; 5070 m) using PacBio HiFi long-read sequencing technology. PacBio HiFi long-read sequences were assembled using hifiasm^28^. To improve chromosome-scale scaffolding of the PacBio HiFi, we also constructed libraries using long-range enzyme-free proximity ligation of endogenous chromatin (Dovetail Omni-C™) followed by deep sequencing with Illumina paired-end (150 bp) technology and assembly using the HiRise™ software^29^. DNA yield from UACH8516 (ethanol-preserved tissue) was of insufficient quantity and quality for scaffolding with the Dovetail Omni-C™ protocol, so we used flash frozen tissue from a second vouchered male specimen (OMNH30107, -34.237974, -69.410897, Mendoza, Argentina) acquired as a loan from the Sam Noble Museum of Natural History, University of Oklahoma. Collectively, we used these data to generate a highly contiguous 2.66 GB assembly (scaffold N50=147.8Mb; N90=59.3Mb; see Figure S3).

### DNA extraction and sequencing

All library preparations for whole genome resequencing experiments were conducted in the University of Montana Genomics Core facility. In the case of live-caught mice, we extracted genomic DNA from ethanol-preserved liver tissue using the DNeasy Blood and Tissue kit or the DNeasy Blood and Tissue QIAcube kit (Qiagen). For the high elevation mummy samples, we extracted DNA from mandible or femur bones in a separate facility following the LK protocol^30^, with modifications to the binding apparatus^31^. Briefly, bones were washed in nuclease free water, followed by agitation in 95% ethanol and then air dried for 1 hour. A 5 mm steel bead was added to the sample and frozen at -80°C for 2 hours. Samples were homogenized on a Tissuelyser (Qiagen) and lysed for 1-3 days at 56°C in buffer ATL and proteinase K with agitation. DNA was isolated following a standard Phenol:Chloroform:Isoamyl Alcohol (24:25:1) protocol and precipitated following a standard NaOAC/Ethanol protocol.

Libraries were prepped using the KAPA HyperPlus kit (Roche). To prepare genomic libraries, contemporary DNA from live-caught mice was sheared on a Covaris E220 sonicator. For the mummy specimens, initial DNA extractions yielded relatively high molecular weight DNA and so we prepared sequencing libraries from sheared DNA as above. We also generated replicate sequencing libraries using unfragmented DNA for five samples. All libraries were prepared with KAPA HiFi DNA Polymerase to help limit DNA misincorporations commonly associated with cytosine deamination of ancient DNA^20^. Individual libraries were indexed using KAPA UDI’s and pooled libraries were sent to Novogene for Illumina sequencing on a Novaseq X.

### Read quality processing and mapping to the reference genome

We used the program fastp 0.23.2^32^ to remove adapter sequences, and to trim and filter low quality reads from sequences generated from fragmented and unfragmented library preps. We used a 5 bp sliding window to remove bases with a mean quality less than 20 and discarded all reads less than 25 bp long with all other settings at default. We merged all overlapping reads that passed filters and retained all reads that could not be merged or whose paired reads failed filtering. We separately mapped merged reads, unmerged but paired reads, and unpaired reads to the reference genome with BWA 0.7.17^33^ using the mem algorithm with the -M option which flags split reads as secondary for downstream compatibility. We sorted, merged, and indexed all resulting binary alignment maps (BAMS) with SAMtools 1.15.1 ^34^. We used picard 2.27.4 to detect and remove PCR duplicates. We used GATK 3.8 ^35^ to perform local realignment around targeted indels to generate our final analysis ready BAM files.

### Sex assignment

We used depth of coverage of the X-chromosome to identify the sex of each sequenced mummy specimen. The same approach correctly identified the sex of all live-trapped mice that were sexed in the field. We used the R package SATC ^36^ to identify sex-linked scaffolds by examining normalized coverage across scaffolds. After identifying scaffold 13 (150.8 Mbp) as X-linked, we assigned sex based on 0.5x (male) or 1.0 x (female) scaffold-specific coverage.

### DNA damage profiles

For genome sequences of the summit mummies, we used the program mapDamage 2.0.9 ^37^ to quantify spatial patterns of DNA base misincorporation along sequencing reads and to quantify the taxonomic identity of contaminant (non-endogenous) DNA. We ran mapDamage with default settings on all aligned reads from all libraries (fragmented and unfragmented) prepared from mummified carcasses and skeletal material.

### Population genomic analysis

We calculated genotype likelihoods for scaffolds 1-19 (covering >90% of the genome) for all samples in ANGSD 0.939 ^38^. For mummy samples where we had sequences from both fragmented and unfragmented libraries, we used alignments from unfragmented libraries. We estimated relationships among samples using NGSrelate 2.0 ^39^ using default settings. We subsampled our dataset and retained a single representative from each identified family/sibling group for all subsequent analyses. We calculated a PCA for our subsampled dataset using PCAngsd 1.10 ^40^. We used NgsAdmix 32 ^41^ to calculate admixture proportions across our subsampled dataset. We ran 10 iterations for *K* values 1-10, where *K* is the number of distinct genetic clusters. To find the estimate of *K* that provided the best fit to our data, we used the Evanno method implemented in the online CLUMPAK ^42^ server. For each value of *K* we merged results across all iterations to represent the average proportion population assignment across runs. All visualizations were generated in R 4.1.3, ggplot2 3.3.6 ^43^, and Pophelper 2.3.1 ^44^.

