## Supplementary Material for "Genomic and radiocarbon insights into the mystery of mouse mummies on the summits of >6000 m Andean volcanoes"

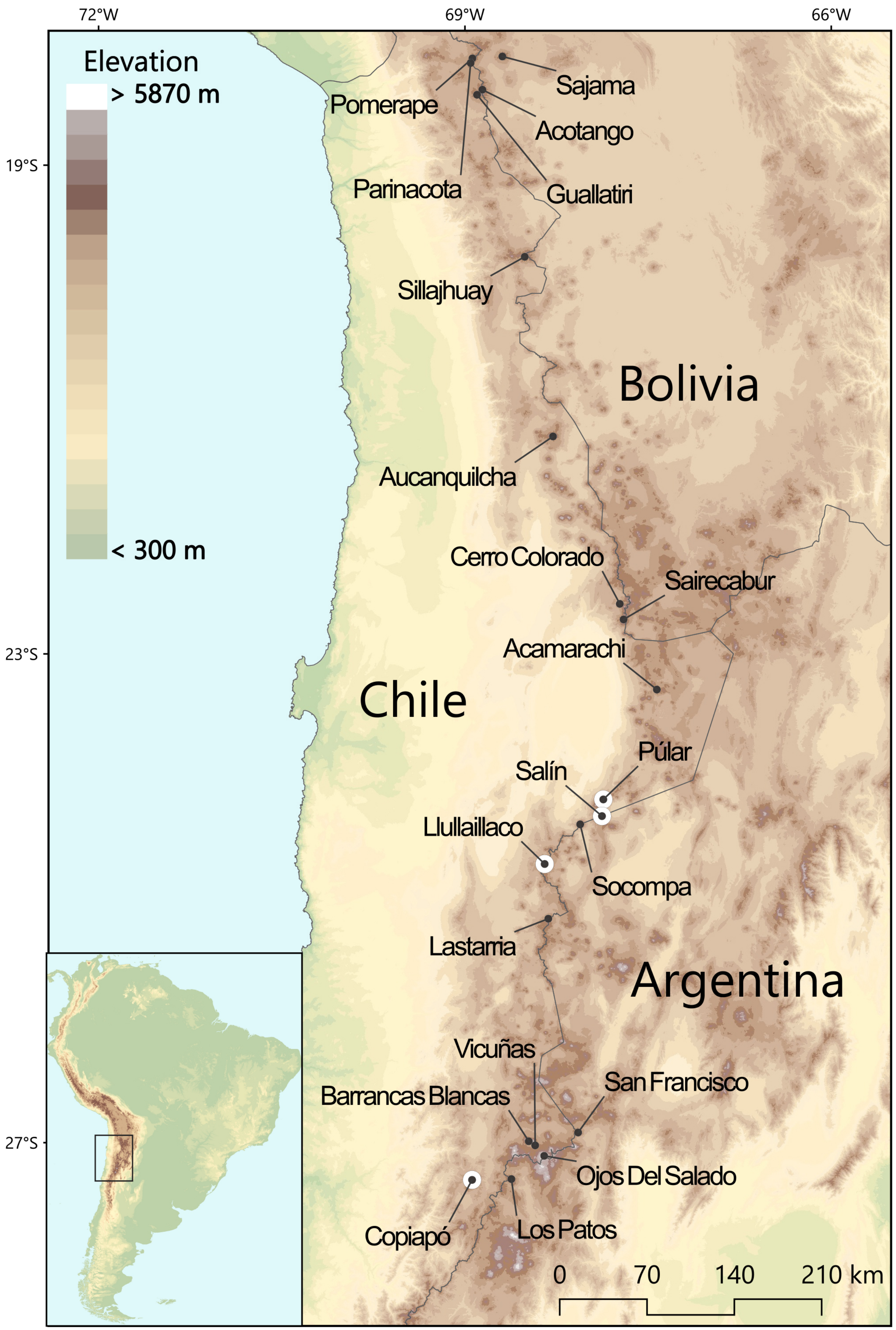


**Figure S1. Surveyed volcanoes in the Central Andes.** White circles denote volcanoes with summit records of *Phyllotis* mice.
From North to South:

| # | Name | Location | Elevation (m) |
|---|---|---|---|
| 1 | Nevado Sajama | Bolivia, División Oruro | 6542 |
| 2 | Volcán Pomerape | Boliva-Chile (División Oruro/Región de Arica y Parinacota) | 6282 |
| 3 | Volcán Parinacota | Boliva-Chile (División Oruro/Región de Arica y Parinacota) | 6319 |
| 4 | Acotango | Boliva-Chile (División Oruro/Región de Arica y Parinacota) | 6052 |
| 5 | Volcán Guallatire | Chile (Región de Arica y Parinacota) | 6071 |
| 6 | Volcán Sillajhuay | Bolivia-Chile (División Oruro/Región de Arica y Parinacota) | 5982 |
| 7 | Volcán Aucanquilcha | Chile (Región de Antofagasta) | 6176 |
| 8 | Cerro Colorado | Chile (Región de Antofagasta) | 5748 |
| 9 | Cerro Sairecabur | Chile (Región de Antofagasta) | 5971 |
| 10 | Volcán Acamarachi | Chile (Región de Antofagasta) | 6046 |
| 11 | Volcán Púlar | Chile (Región de Antofagasta) | 6233 |
| 12 | Volcán Salín | Argentina-Chile (Provincia de Salta/Región de Antofagasta) | 6029 |
| 13 | Volcán Socompa | Argentina-Chile (Provincia de Salta/Región de Antofagasta) | 6051 |
| 14 | Volcán Llullaillaco | Argentina-Chile (Provincia de Salta/Región de Antofagasta) | 6739 |
| 15 | Volcán Lastarria | Argentina-Chile (Provincia de Salta/Región de Antofagasta) | 5706 |
| 16 | Volcán San Francisco | Argentina-Chile (Provincia de Catamarca/Región de Atacama) | 6016 |
| 17 | Barrancas Blancas | Chile (Región de Atacama | 6119 |
| 18 | Cerro Vicuñas | Chile (Región de Atacama | 6067 |
| 19 | Nevado Ojos del Salado | Argentina-Chile (Provincia de Catamarca/Región de Atacama) | 6893 |
| 20 | Volcán Los Patos | Argentina-Chile (Provincia de Catamarca/Región de Atacama) | 6239 |
| 21 | Volcán Copiapó | Chile (Región de Atacama) | 6052 |

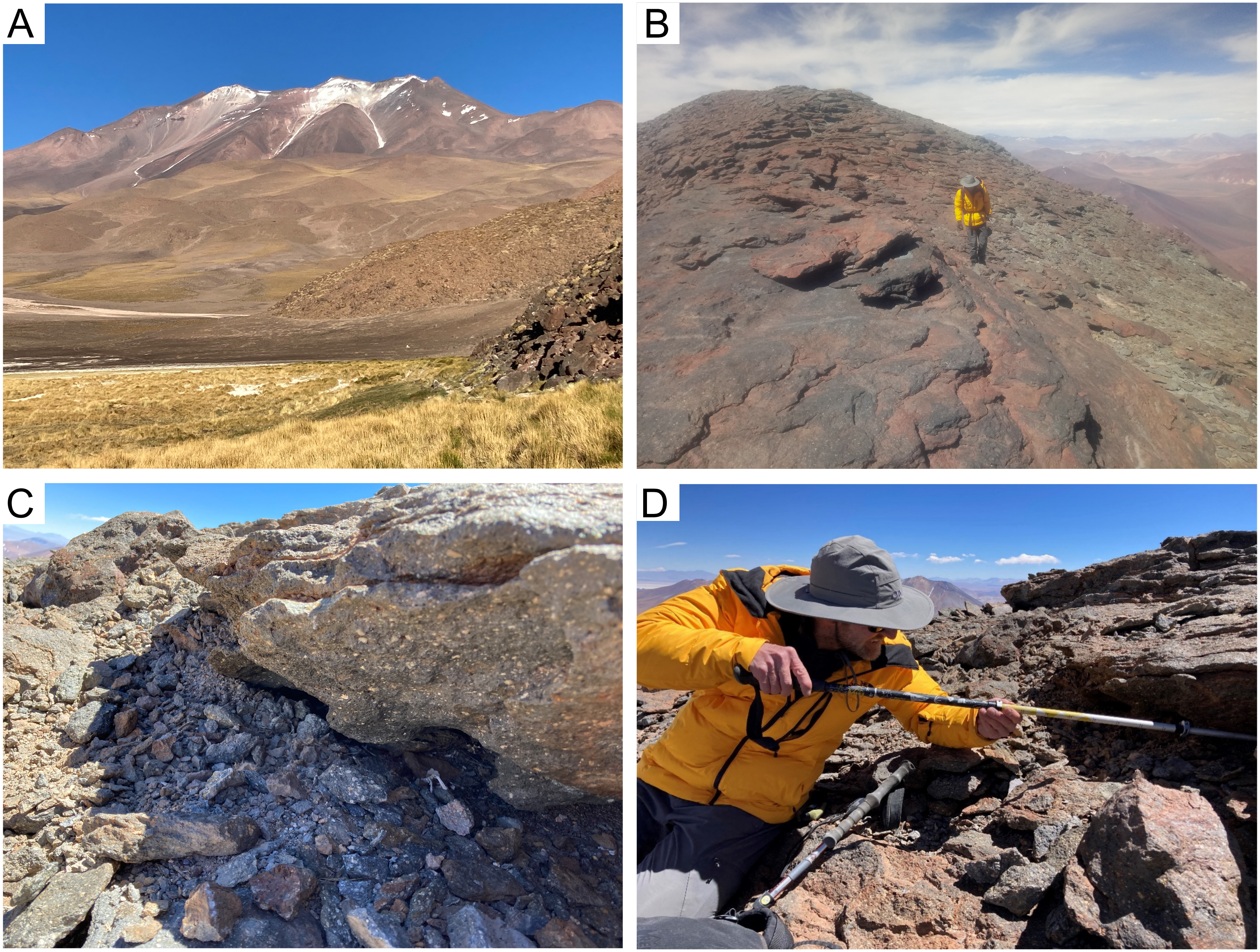

**Figure S2. Discovery of mouse graveyard on the summit of Volcán Púlar.** (A) View of Volcán Púlar from the southeast. (B) View of exposed volcanic rock on the summit of Volcán Púlar (6233 m). (C) Rocky alcove on the summit of Volcán Púlar where we excavated one individually intact *Phyllotis* mummy (UACH8537), along with partial skeletal remains of three other individual mice (UACH8546-8548). (D) Excavation of skeletal remains from beneath the rock ledge.

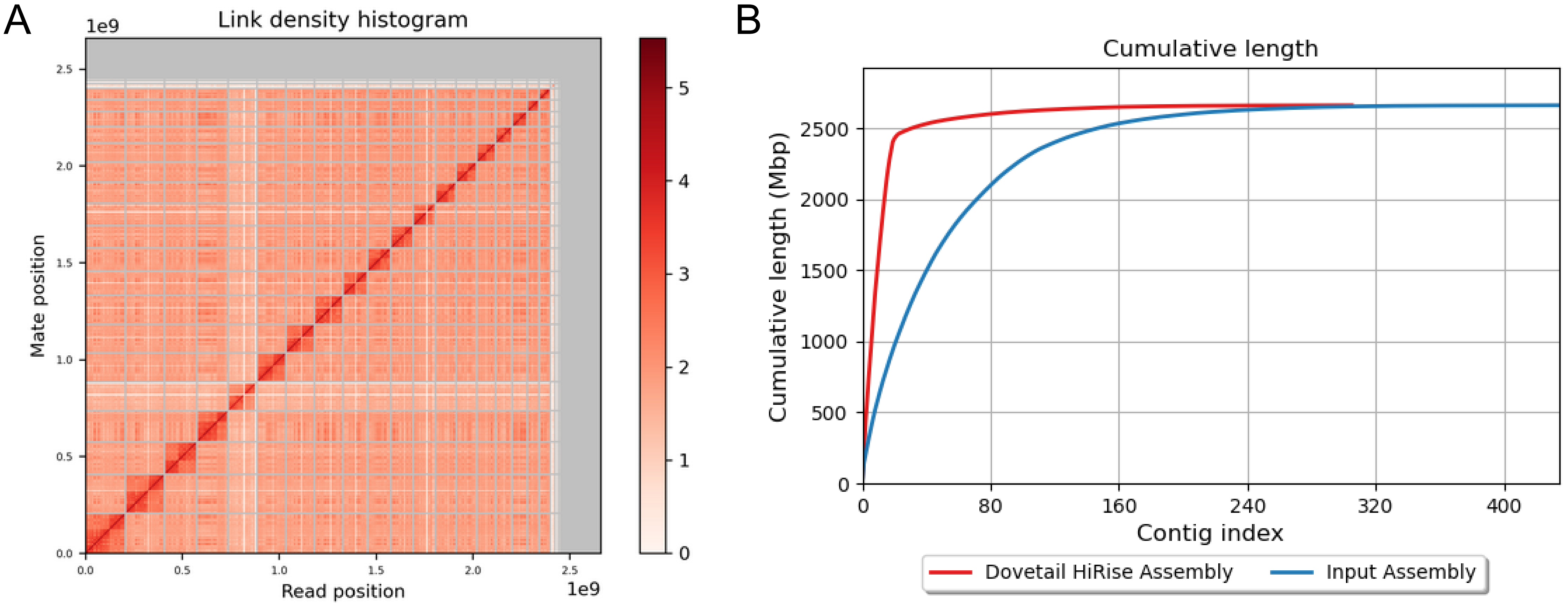


**Figure S3. Scaffolding-plot of *Phyllotis vaccarum* genome assembly.** (A) Binned mapping positions of first and second Omni-C read pairs, with scaffolds ordered by size and bordered by gray lines. (B) Assembly improvement curve showing reduction in L90 with inclusion of OmniC linked reads (HiRise Assembly) compared to PacBio sequencing only (input assembly).

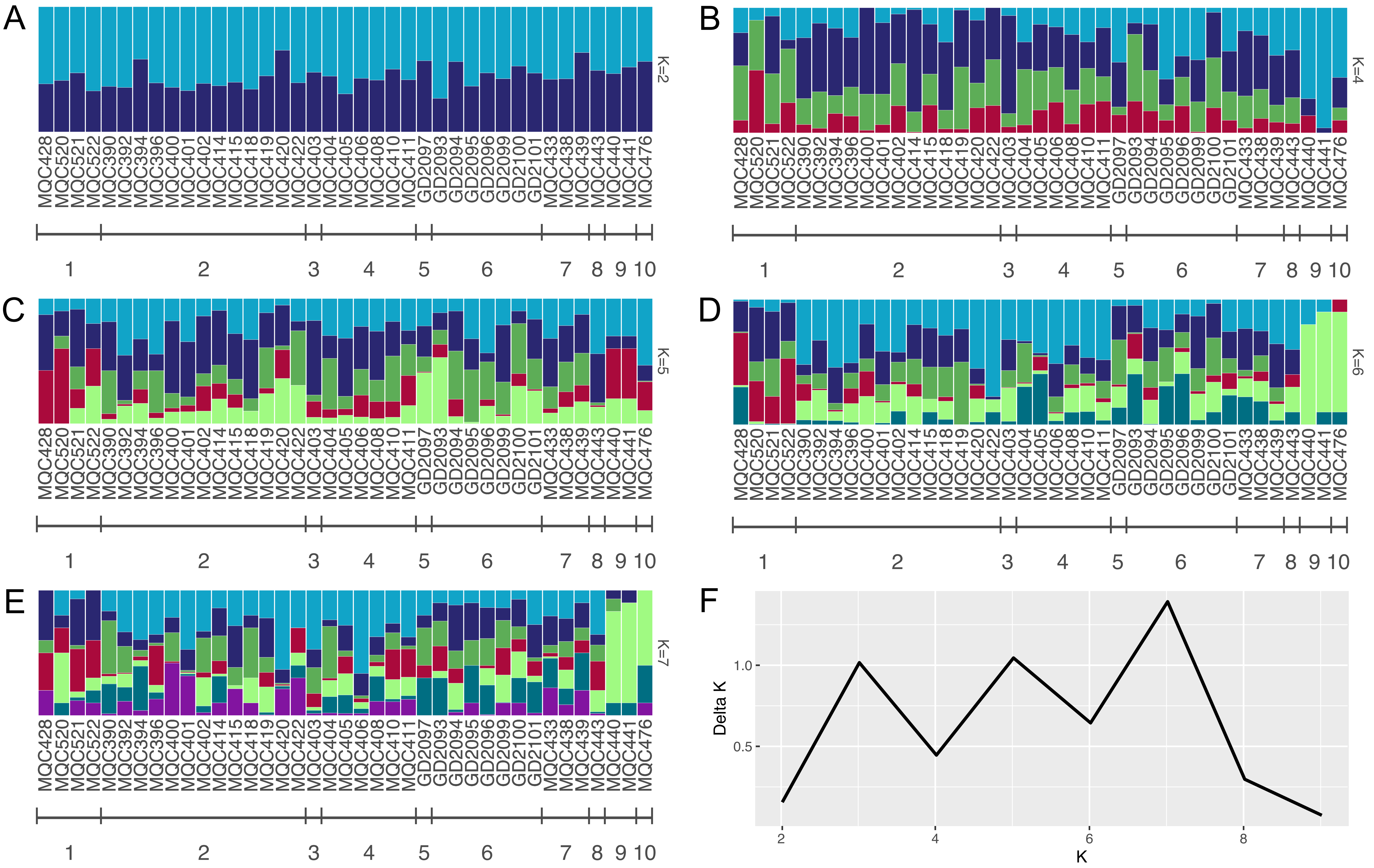


**Figure S4. Low population structure across the surveyed region, as revealed by a model-based clustering analysis of genomic polymorphism data.** Admixture plots for K=2, 4-7 genetic clusters show that proportional assignments of ancestry were simila for mice from all localities (admixture plot for K=3 is shown in Figure 3B). Delta K, calculated using the Evanno method, shows multiple peaks at K=3, 5, and 7.

| UACH ID# | Total reads | Mapped reads | Mean coverage | Percent aligned | Total reads (unfragmented) | Mapped reads (unfragmented) | Mean coverage (unfragmented) | Percent aligned (unfragmented) |
|---|---|---|---|---|---|---|---|---|
| 8538 | 35270349 | 35093894 | 1.93x | 99.50% | | | | |
| 8539 | 32421746 | 32291962 | 1.76x | 99.60% | | | | |
| 8540 | 28912450 | 28790027 | 1.55x | 99.58% | | | | |
| 8541 | 15171858 | 15024039 | 0.83x | 99.03% | | | | |
| 8542 | 30823587 | 30695265 | 1.59x | 99.58% | | | | |
| 8543 | 37473122 | 37317683 | 1.99x | 99.59% | | | | |
| 8544 | 27725041 | 27647373 | 1.62x | 99.72% | | | | |
| 8545 | 45103193 | 44842990 | 2.56x | 99.42% | | | | |
| 8555 | 18489767 | 14150982 | 0.63x | 76.53% | 266833096 | 148477017 | 7.66x | 55.64% |
| 8537 | 280708204 | 183987607 | 9.09x | 65.54% | 120863677 | 102494211 | 5.89x | 84.80% |
| 8546 | 240533459 | 196126272 | 11.13x | 81.54% | 268916882 | 230888539 | 13.24x | 85.86% |
| 8547 | 145609511 | 37886116 | 1.79x | 26.02% | 144433586 | 58707035 | 3.15x | 40.65% |
| 8548 | 168632141 | 91343188 | 4.94x | 54.17% | 130679148 | 98430814 | 5.73x | 75.32% |

**Table S1**. **Sequencing results for summit mummies**. Results from low-coverage whole genome resequencing of mummified mouse cadavers and skeletal remains. For samples collected from the summits of Volcán Copiapó (UA8555) and Volcán Púlar (UACH8537, 8546, 8547, and 8548), we prepared both fragmented and unfragmented genomic libraries.
